# Kv2/Kv6.4 heteromeric potassium channels are expressed in spinal motor neurons and localized at C-bouton synapses

**DOI:** 10.1101/2025.06.04.657913

**Authors:** Taylor A. Lacey, Karl D. Murray, James S. Trimmer, Jon T. Sack, Michael J. Ferns

## Abstract

Voltage-gated K^+^ channels of the Kv2 family co-assemble with electrically silent KvS subunits in specific subpopulations of brain neurons, forming heteromeric Kv2/KvS channels with distinct functional properties. Little is known about the composition and function of Kv2 channels in spinal cord neurons, however. Here, we show that while Kv2.1 is broadly expressed in multiple classes of spinal cord neurons, the Kv6.4 “electrically-silent” subunit is specifically expressed in motor neurons. In motor neurons, we find that Kv6.4 protein is co-clustered with Kv2.1 and Kv2.2 subunits at endoplasmic reticulum-plasma membrane (ER-PM) junctions beneath C-bouton synapses. In Kv2.1 S590A mutant mice, in which Kv2.1 is unable to bind ER VAP proteins, Kv2.1 and Kv6.4 clustering at ER-PM junctions is severely reduced suggesting Kv2 channels are localized at ER-PM junctions by the same molecular mechanism in motor neurons and brain neurons. Moreover, clustering of Kv6.4, as well as the AMIGO-1 auxiliary subunit, are severely reduced in Kv2.1 knockout mice and moderately reduced in Kv2.2 knockout mice. Thus, expression and localization of Kv6.4 subunits is dependent on Kv2 subunits, likely through their co-assembly into heteromeric channels. Finally, we find that presynaptic C-boutons and postsynaptic clusters of the ER-resident sigma1-receptor are preserved in motor neurons of Kv2 knockout mice. Together, these findings identify a specific Kv2/KvS channel subtype expressed in motor neurons that localizes to C-bouton junctions where it could regulate neuronal excitability and signaling at ER-PM junctions.

## 1. Introduction

The Kv2 family of voltage-gated K^+^ channels consists of two α subunits, Kv2.1 and Kv2.2, which form homo- and hetero-tetrameric channels that are robustly expressed throughout the nervous system (Bishop et al., 2015). Kv2 channels play an important role in regulating neuronal action potentials and excitability in many brain neurons (Guan et al., 2007; McKeown et al., 2008; Liu and Bean, 2014; Trimmer, 2015). Indeed, Kv2.1 knockout (KO) mice have enhanced neuronal activity and are susceptible to seizures (Speca et al., 2014), and patients with de novo mutations in Kv2.1 have epileptic encephalopathy (Allen et al., 2020). In addition, Kv2 channels play a separate structural role in organizing endoplasmic reticulum - plasma membrane (ER-PM) junctions on neuronal cell bodies and proximal dendrites (Fox et al., 2015; Kirmiz et al., 2018b). Kv2 channels create these junctions by binding VAP proteins in the underlying ER in a phosphorylation dependent manner (Lim et al., 2000; Johnson et al., 2018; Kirmiz et al., 2018a), and phosphorylation at these sites is regulated by electrical activity and other physiological stimuli (Misonou et al., 2004; Misonou et al., 2005; Misonou et al., 2006). Numerous signaling proteins have been shown to be recruited to Kv2-containing ER-PM junctions, including L-type Ca^2+^ channels (LTCCs) and an array of Ca^2+^-regulated signaling proteins (Vierra et al., 2019), lipid-handling proteins (Kirmiz et al., 2019; Sun et al., 2019), and protein kinase A (PKA) signaling proteins (Vierra et al., 2023). Thus, these microdomains are thought to serve as important hubs for somatodendritic Ca^2+^, lipid, and PKA signaling in brain neurons (Johnson et al., 2019; Vierra and Trimmer, 2022; Guillen-Samander and De Camilli, 2023; Vierra, 2024). The AMIGO-1 auxiliary subunit is also localized at ER-PM junctions through its association with Kv2 channels and modulates Kv2 channel gating (Bishop et al., 2018; Maverick et al., 2021; Sepela et al., 2022).

Another unusual feature of the Kv2 family is that they co-assemble with the distinct KvS family of voltage-gated K^+^ channel subunits to form heteromeric channels. The KvS subunit family consists of 10 members (Kv5.1, Kv6.1-6.4, Kv8.1-8.2, Kv9.1-9.3) which share homology with Kv2 subunits and are well conserved among diverse vertebrate species (Bocksteins and Snyders, 2012; Bocksteins, 2016). KvS subunits were originally designated as “electrically silent” as they do not form functional channels when expressed alone in heterologous cells. However, multiple studies demonstrate that they co-assemble selectively with Kv2 family members to form heteromeric KvS/Kv2 channels with biophysical properties distinct from those of Kv2 homomers (Bocksteins and Snyders, 2012). Moreover, while Kv2 subunits are broadly expressed, KvS subunit transcripts are often expressed in restricted regions and neuronal subpopulations in the CNS (Castellano et al., 1997; Salinas et al., 1997; Kramer et al., 1998). Consequently, distinct Kv2/KvS channel subtypes may be expressed in different neurons, where they may serve neuron-specific functions. For this reason, KvS subunits have considerable potential as drug targets, as specific Kv2/KvS channel modulators could regulate neuronal activity with a higher degree of specificity than Kv2 modulators. Consistent with this, we recently showed that within brain, the Kv5.1 subunit is selectively expressed in specific layers of neocortex, where it is localized at ER-PM junctions on the soma and proximal dendrites of neurons (Ferns et al., 2025). The localization and potential function of other KvS subunits in CNS neurons remains largely unknown.

In spinal cord, Kv2 channels are expressed in many neuronal subtypes but are particularly prominent in motor neurons, where their precise function remains poorly understood. Some studies have shown that Kv2 channels regulate excitability and firing of neonatal motor neurons in a complex manner that is dependent on patterns of stimulation (Romer et al., 2019; Deardorff et al., 2021). A recent study, however, found Kv2 channels have minimal impact on excitability of more mature motor neurons (Smith et al., 2024). Kv2.1 channels could also play a structural role as they form large (micron size) clusters on the motor neuron cell bodies and proximal dendrites immediately beneath cholinergic C-bouton synapses (Muennich and Fyffe, 2004; Wilson et al., 2004). C-bouton synapses arise from Pitx2-positive cholinergic interneurons in spinal cord (Zagoraiou et al., 2009), and stimulation of these inputs is thought to facilitate repetitive firing of motor neurons by reducing spike half-width and enhancing afterhyperpolarizations (Nascimento et al., 2020). Intriguingly, C-bouton synapses are associated with ER-PM junctions and several proteins localize to these sites, including the muscarinic m2 receptor (Wilson et al., 2004), SK channels (Deardorff et al., 2013), the sigma-1 receptor (S1R) (Mavlyutov et al., 2010; Mavlyutov et al., 2012), and neuregulin (Gallart-Palau et al., 2014; Casanovas et al., 2017). Calcium and lipid signaling components have not been detected at C-bouton associated ER-PM junctions on motor neurons, and consequently they likely differ functionally from ER-PM junctions in brain neurons. Importantly, C-bouton junctions appear critical for motor neuron homeostasis and survival, as mutations in VAPB (Nishimura et al., 2004; Borgese et al., 2021), S1R (Luty et al., 2010; Al-Saif et al., 2011), and neuregulin/ErbB4 signaling (Takahashi et al., 2013) have been identified as familial causes of amyotropic lateral sclerosis (ALS). Perturbations in C-bouton junctions have also been observed in sporadic and other familial forms of ALS, as well as following nerve injury (Gallart-Palau et al., 2014; Mavlyutov et al., 2015; Casanovas et al., 2017).

To gain insight into the role of Kv2/KvS channel complexes in spinal cord, we investigated the composition and function of Kv2 channels at C-bouton synapses on motor neurons. We show that a KvS subunit family member, Kv6.4, is specifically expressed in motor neurons and is co-clustered with Kv2 subunits at C-bouton synapses. We further show that Kv6.4 clustering at C-bouton synapses is significantly decreased in gene-edited mice harboring the Kv2.1 S590A mutation that is unable to bind ER VAP proteins, as well as in Kv2.1 and Kv2.2 KO mice. These findings provide evidence that Kv2/Kv6.4 heteromeric channels are selectively expressed in motor neurons, where they could modulate neuronal excitability and signaling at ER-PM junctions.

## 2. Materials and Methods

### DNA constructs

Plasmids encoding untagged and GFP- or HA-tagged rat Kv2.1 and Kv2.2 have been previously described (Lim et al., 2000; Bishop et al., 2018; Kirmiz et al., 2018a). Plasmids encoding mouse KvS subunits with C-terminal myc-DDK tags were obtained from Origene (Kv6.1 (MR223857); Kv6.4 (MR224440); Kv9.1 (MR218800) and Kv9.3 (MR217315). A plasmid encoding human Kv6.4 with an N-terminal HA epitope tag was generated by VectorBuilder (en.vectorbuilder.com).

### Animals

All procedures using mice and rats were approved by the University of California, Davis Institutional Animal Care and Use Committee and performed in accordance with the NIH Guide for the Care and Use of Laboratory Animals. Animals were maintained under standard light-dark cycles and allowed to feed and drink ad libitum. Adult C57BL/6J mice (RRID:IMSR_JAX:000664) 12-16 weeks old of both sexes were used in immunolabeling experiments. Kv2.1-KO mice (RRID:IMSR_MGI:3806050) have been described previously (Jacobson et al., 2007; Speca et al., 2014), and were generated from breeding of Kv2.1^+/-^ mice that had been backcrossed on a C57/BL6J background. Kv2.2-KO mice have been described previously (Hermanstyne et al., 2010; Hermanstyne et al., 2013), and Kv2.1 and Kv2.2 double-KO (Kv2 dKO) mice (Kv2.1^−/−^/Kv2.2^−/−^) were generated by breeding Kv2.1^+/−^ mice with Kv2.2^−/−^ mice. Kv2.1 S590A gene edited mice were generated by CRISPR/Cas-9 gene editing and have been described previously (Matsumoto et al., 2023).

### Antibodies

All primary antibodies used in this study are listed in Table 1. Monoclonal antibodies were generated against mouse Kv6.4 by the UC Davis/NIH NeuroMab facility (https://neuromab.ucdavis.edu). Positive hybridoma subclones identified in the initial screen were re-tested in multiple applications, and clone N458/10 was found to recognize Kv6.4 by immunolabeling of transfected HEK293T (HEK) cells, immunoprecipitations (IPs), immunoblotting of transfected cell lysates, and immunolabeling of spinal cord tissue sections. Similar applications were used to test the specificity of N458/10 for Kv6.4 compared to Kv2.1, Kv2.2, and other KvS subunits. Characterization of the anti-Kv6.4 mAb N458/10 is detailed in Figure 1.

**Figure 1.**
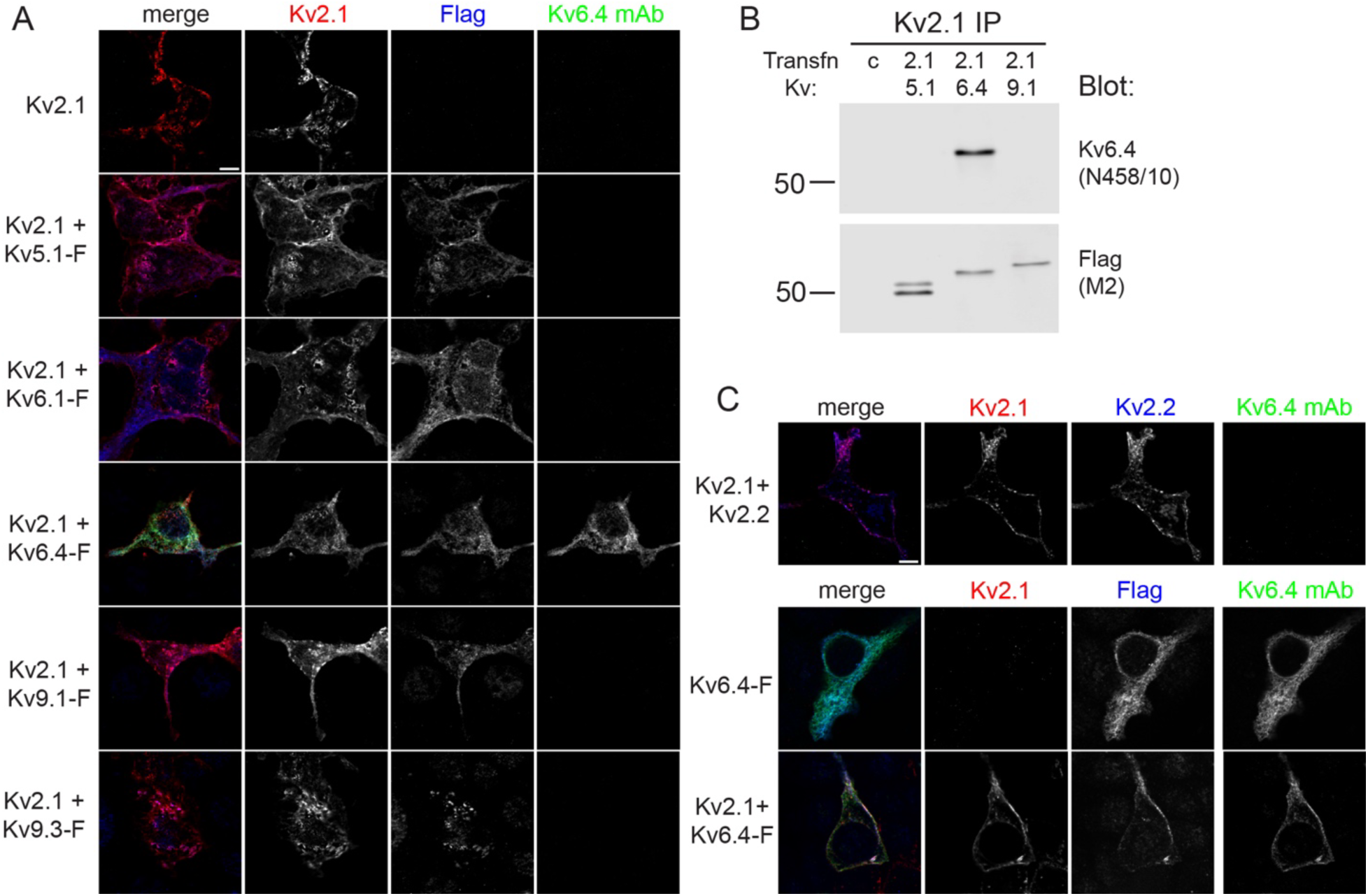
Characterization of the anti-Kv6.4 mAb N458/10. (A) HEK cells expressing Kv2.1 alone or Kv2.1 plus Flag-tagged KvS subunits were fixed in 4% formaldehyde in PBS, and immunolabeled with antibodies against Kv2.1 (K89/34, red), Flag (pAb, blue) and Kv6.4 (N458/10, green). The N458/10 mAb recognized Kv6.4 but not Kv2.1 or other KvS subunits. Scale bar = 10 μm. (B) Kv2/KvS channels were immunoprecipitated from lysates of HEK cells expressing Kv2.1 plus Flag-tagged-Kv5.1, Kv6.4 or Kv9.1 subunits using an anti-Kv2.1 antibody. The products were size fractionated on SDS gels and immunoblotted for Kv6.4 using N458/10. N458/10 detected a single band for Kv6.4 at the expected molecular weight of ∼57 kD and did not recognize the immunoprecipitated Kv5.1 or Kv9.1. Immunoblotting with anti-Flag antibody confirmed the presence of all three KvS subunits in the immunoprecipitation products. Numbers to the left are molecular weights standards in kD. (C) HEK cells expressing Kv2.1 and Kv2.2, Kv6.4 alone, or Kv2.1 and Kv6.4 were fixed in 4.5% glyoxal/4% acetic acid for 15 min, and immunolabeled with antibodies against Kv2.1 (K89/34, red), Kv2.2 (2.2C pAb, blue), and Kv6.4 (N458/10, green), or Kv2.1 (K89/34, red), Flag (pAb, blue), and Kv6.4 (N458/10, green). Using these fixation conditions, N458/10 antibody detected Kv6.4 on the cell periphery when it was co-expressed with Kv2.1. N458/10 did not recognize Kv2.1 or Kv2.2. Scale bar = 10 μm.

**Table 1.** Antibody Information.

| **Antigen and Antibody** | **Immunogen** | **Manufacturer information** | **Concentration used** |
| --- | --- | --- | --- |
| **Kv2.1 (KC)** | Synthetic peptide aa 837-853 of rat Kv2.1 | Rabbit pAb, In-house (Trimmer Laboratory), RRID:AB_2315767 | Affinity purified, 1:100 |
| **Kv2.1 (K89/34)** | Synthetic peptide aa 837-853 of rat Kv2.1 | Mouse IgG1 mAb, NeuroMab, RRID: AB_2877280 | Purified, 1-2 µg/ml |
| **Kv2.1 (K89/34R)** | Synthetic peptide aa 837-853 of rat Kv2.1 | Mouse IgG2b mAb, In-house (Trimmer Laboratory), RRID: AB_2934194 | Tissue culture supernatant, 1:10 |
| **Kv2.2 (Kv2.2C)** | Recombinant protein aa 717-907 of rat Kv2.2 | Rabbit pAb, in-house (Trimmer Laboratory), RRID:AB_2801484 | Affinity purified, 1:100 |
| **Kv2.2 (N372B/60)** | Recombinant protein aa 717-907 of rat Kv2.2 | Mouse IgG2a mAb, NeuroMab, RRID: AB_2877437 | Purified, 10 µg/mL |
| **Kv6.4 (N458/10)** | Recombinant protein aa 424-494 of mouse Kv5.1 | Mouse IgG2a mAb, NeuroMab, RRID:AB_2877578 | Tissue culture supernatant, 1:10<br>8 µg/mL |
| **Kv2.1 phospho-S590 (L100/1)** | Synthetic phosphopeptide aa 581-590 of rat Kv2.1 | Mouse IgG1 mAb, In-house (Trimmer Laboratory), RRID: AB_2531886 | Purified, 5 µg/ml |
| **Ankyrin G**<br>**(N106/36)** | Full-length<br>recombinant human<br>Ankyrin G | Mouse IgG2a mAb, NeuroMab,<br>RRID:AB_2877524 | Tissue culture<br>supernatant, 1:5 |
| **AMIGO-1**<br>**(L86A/37)** | Recombinant protein<br>aa 395-493 of mouse<br>AMIGO-1 | Mouse IgG2b mAb, NeuroMab,<br>RRID:AB_2877364 | Tissue culture<br>supernatant, 1:5 |
| **ChAT**<br>**(1E6)** | ChAT purified from<br>rat brain | Mouse IgG1 mAb,<br>Millipore #MAB305,<br>RRID:AB_94647 | Ascites fluid, 1:200 |
| **ChAT** | Synthetic peptide aa<br>629-642 of mouse<br>ChAT | Goat pAb,<br>Novus #NB100-40794,<br>RRID:AB_714957 | Affinity-purified,<br>2 µg/ml |
| **Grp75/Mortalin**<br>**(N52A/42)** | n/a | Mouse IgG1 mAb, NeuroMab,<br>RRID:AB_2877320 | Tissue culture<br>supernatant, 1:10 |
| **Flag** | Synthetic peptide,<br>DYKDDDDK | Rabbit pAb,<br>Rockland #600-401-383,<br>RRID:AB_219374 | Affinity purified,<br>1.2 µg/ml |
| **GFP** | Recombinant<br>protein | Rabbit pAb,<br>Rockland #600-401-215<br>RRID:AB_828167 | Affinity purified,<br>1.25 µg/ml |
| **HA**<br>**(12CA5)** | Human influenza<br>virus hemagglutinin<br>protein aa 98-106 | Mouse IgG2b mAb,<br>RRID:AB_2532070 | Purified,<br>5 µg/mL |
| **Sigma1-R**<br>**(B-5)** | Human sigma-1-receptor aa 136-169 | Mouse IgG1 mAb, Santa Cruz<br>Biotechnology # sc-137075,<br>RRID: AB_2285870 | Purified,<br>1:400 |
| **Synaptophysin**<br>**(EP1098Y)** | Unspecified | Rabbit mAb<br>Epitomics #1870,<br>RRID: AB_882786 | Purified,<br>1:200 |

### HEK293T cell culture and transfection

HEK293T cells (ATCC Cat# CRL-3216) were maintained in Dulbecco’s modified Eagle’s medium (Gibco Cat# 11995065) supplemented with 10% Fetal Clone III (HyClone Cat# SH30109.03), 1% penicillin/ streptomycin, and 1x GlutaMAX (ThermoFisher Cat# 35050061) in a humidified incubator at 37 °C and 5% CO_2_. Cells were transiently transfected using Lipofectamine 3000 (Life Technologies, cat. #L3000008) following the manufacturer’s protocol, then rescued in fresh growth medium, and used for experiments 36-48 hours after transfection. For immunolabeling, cells were plated and transfected on poly-L-lysine (Sigma Cat# P1524)-coated microscope cover glasses (VWR Cat# 48366-227). For immunoblotting, cells were grown and transfected on 60 mm tissue culture dishes.

### Immunolabeling of HEK cells

HEK293T cells were fixed for 15 mins at 4 °C either in 4% formaldehyde prepared fresh from paraformaldehyde in PBS buffer pH 7.4, or 4.5% glyoxal/4% acetic acid at 20 °C (Richter et al., 2018; Konno et al., 2023). All subsequent steps were performed at RT. Cells were then washed 3 x 5 mins in PBS, followed by blocking in blotto-PBS (phosphate-buffered saline supplemented with 4% (w/v) non-fat milk powder and 0.1 % (v/v) Triton X-100 [Roche Cat# 10789704001]) for 1 hour. Cells were immunolabeled for 1-2 hours with primary antibodies diluted in blotto-PBS and subsequently washed 3 x 5 mins in blotto-PBS. They were then incubated with mouse IgG subclass- and/or species-specific Alexa-conjugated fluorescent secondary antibodies (Invitrogen) diluted in blotto-PBS for 45 min and washed 3 x 5 mins in PBS. Cover glasses were mounted on microscope slides with Prolong Gold mounting medium (ThermoFisher Cat # P36930) according to the manufacturer’s instructions.

### Immunolabeling of spinal cord sections

Following administration of sodium pentobarbital (Nembutal, 60 mg/kg) to induce deep anesthesia, mice were transcardially perfused with either 4% formaldehyde (freshly prepared from paraformaldehyde) in 0.1 M sodium phosphate buffer pH 7.4, or 2% formaldehyde (prepared fresh from paraformaldehyde) in 0.05 M Na acetate buffer pH 6.0 (Berod et al., 1981), or 9% glyoxal/8% acetic acid (Richter et al., 2018; Konno et al., 2023). Fixed spinal cords were removed from the vertebral column and embedded in OCT blocks for frozen sectioning. Spinal cord sections (12-14 µm thick) were cut on a cryostat, placed on slides, and permeabilized and blocked in PBS containing 10% goat serum and 0.3% Triton X-100 for 1 hour at RT. The sections were then incubated with primary antibodies (Table 1) diluted in PBS containing 5% goat serum and 0.1% Triton X-100, either for 3 hr at room temperature or overnight at 4 °C. After 4 x 5 min washes in PBS plus 0.1% Triton X-100, sections were incubated with goat anti-mouse IgG subclass- and/or species-specific Alexa-conjugated fluorescent secondary antibodies (Invitrogen) for 1 hour. For immunolabeling experiments using goat anti-ChAT polyclonal antibody, sections were blocked in PBS containing 5% donkey serum and primary antibodies detected using Alexa-conjugated fluorescent donkey anti-goat and donkey anti-mouse secondary antibodies. After 3 x 5 min washes in PBS, the sections were coverslipped using Prolong Gold (ThermoFisher Cat # P36930). Immunolabeling using anti-Kv6.4 (N458/10) antibody required antigen retrieval steps for some fixation conditions. For spinal cords fixed with 4% formaldehyde, sections were pre-treated with protease IV as detailed in the RNAscope protocol below (Advanced Cell Diagnostics). For spinal cords fixed with 2% formaldehyde, pH 6, sections were pre-treated with 0.1% SDS for 5 min. Sections fixed with Glyoxal did not require antigen retrieval.

### RNAscope *in situ* hybridization

Cryostat sections (15 µm) were cut from lumbar spinal cord of adult mice which had been perfusion-fixed with 4% formaldehyde (from paraformaldehyde) in 0.1 M sodium phosphate buffer pH 7.4, as described above. Sections were processed for RNA *in situ* detection using a slightly modified version (Griffith et al., 2019) of the manufacturer’s RNAscope protocol (Advanced Cell Diagnostics) which includes a protease antigen retrieval step. Following *in situ* hybridization with *Kcng4* (Kv6.4) probe (ACD Cat # 316931), sections were blocked with 5% normal goat serum in PBS plus 0.1% Triton X-100) and incubated with anti-Kv2.1 (K89/34) and anti-Kv6.4 (N458/10) mAbs overnight at 4 °C (see Table 1 for details). Sections were washed in PBS, incubated with secondary antibodies for 1 hr at RT, washed again in PBS, and mounted with Prolong-Gold.

### Image acquisition and analysis

Widefield fluorescence images were acquired with an AxioCam MRm digital camera installed on a Zeiss AxioImager M2 microscope or with an AxioCam HRm digital camera installed on a Zeiss AxioObserver Z1 microscope with a 63×/1.40 NA Plan-Apochromat oil immersion objective and an ApoTome coupled to Axiovision software version 4.8.2.0 (Zeiss, Oberkochen, Germany). Motor neurons were distinguished from interneurons by soma size, morphology, laminar location, and their prominent, micron-sized Kv2.1 clusters. Morphological and colocalization analyses of immunolabeled proteins were performed using Fiji software. For analysis of Kv2.1, Kv2.2 and Kv6.4 colocalization, images were subjected to “rolling ball” background subtraction and an ROI was drawn around each motor neuron soma (excluding any areas with dense, autofluorescent Nissl bodies). Pearson’s correlation coefficient (PCC) values were then collected using the Coloc2 plugin. Relative clustering of Kv2 and Kv6.4 was measured by quantifying the coefficient of variation (CV) of the fluorescence intensity (s.d./mean of pixel intensity) in each cell, with a greater CV indicative of a higher degree of protein clustering (Cobb et al., 2015). For comparison of wild-type (WT) and Kv2.1 and Kv2.2 KO samples, analysis was performed on images of spinal cord sections taken using the same exposure times. For this, ROIs consisted of a 2 µm band encompassing the cell surface of each motor neuron soma. Overlapping synaptophysin, AMIGO-1 and S1R puncta in WT and Kv2.1 KOs were quantified using the SynapseCounter plugin. For presentation, images were linearly scaled for min/max intensity in Fiji and saved as RGB TIFFs.

### Immunoprecipitations and Immunoblotting

For immunoblot analyses of transfected HEK cells, cells were solubilized in PBS containing 0.5% Triton X-100, 1 mM PMSF and a protease inhibitor cocktail (Cell Signaling, Cat # 5871). Insoluble material was pelleted by centrifugation at 12,000 x g for 10 min at 4 °C and a sample of the lysate was removed and incubated with 2x SDS loading buffer at 90 °C for 3 min. For IPs, antibodies were added to the remaining supernatant and incubated for 2 hr at 4 °C on a rotator or rocking platform, and then magnetic protein G beads (ThermoFisher Cat# 10004D) were added for an additional 1 hr. After washing 3x in PBS containing 0.5%Triton X100 and protease inhibitors, the IPed proteins were eluted in 2x SDS loading buffer at 80 °C for 4 min. All samples were separated on 7-8% SDS-PAGE gels and transferred to PVDF membrane. Immunoblots were blocked in LiCor Intercept PBS blocking buffer (LiCor Biosciences) and probed for Kv6.4 (N458/10 mAb) or for the relevant GFP or Flag epitope tag. Grp75/Mortalin (mAb N52A/42) was used as a loading control. Dye-conjugated fluorescent secondary antibodies (LiCor Biosciences) were used to detect bound primary antibodies and imaged using an Odyessy DLx scanner (LiCor Biosciences). A dye-conjugated light chain-specific anti-rabbit secondary antibody (Jackson Labs) was used to detect Flag-tagged KvS subunits, to avoid recognition of rabbit antibody heavy chain used for the IPs. Immunoblots were analyzed using Image Studio software (LiCor Biosciences) and statistical analysis was performed using GraphPad Prism (GraphPad.com).

### Statistical analysis

Measurements derived from immunoblots and image analysis were imported into GraphPad Prism for statistical analysis and presentation. Reported values are mean ± s.e.m. We used a one-way ANOVA to compare multiple experimental groups, with post-hoc Tukey’s or Sidak’s multiple comparisons tests to determine which individual means differed. A one sample t and Wilcoxon test was used to compare experimental groups to a normalized, control group. For all tests, P values **<**0.05 were considered to be significantly different, and exact *P*-values are reported in each figure or figure legend.

## 3. Results

### Kv6.4 is expressed in motor neurons

Gene expression studies (Muller et al., 2014) and publicly available databases (*e.g*., Allen Spinal Cord Atlas) identify *Kcng4* (encoding Kv6.4) as a KvS subunit whose transcripts are present in mouse spinal cord including potentially being enriched in motor neurons. To begin to characterize Kv6.4 expression in spinal cord at the protein level, we screened several monoclonal antibodies generated against Kv6.4 by the UC Davis/NIH NeuroMab facility. We found that among these N458/10 recognized mouse Kv6.4 expressed in heterologous cells by immunofluorescence (IF) labeling and immunoblotting. Moreover, we confirmed that in IF experiments N458/10 does not cross-react to Kv2.1, Kv2.2 or to several other KvS subunits expressed in heterologous cells (Figure 1A). However, we found that in cells fixed with 4% formaldehyde, N458/10 failed to detect Kv2/Kv6.4 channels expressed on the plasma membrane of heterologous cells, even though Kv6.4 was co-immunoprecipitated with Kv2.1 from cell lysates (Figure 1B), and Kv2.1/Kv6.4 heteromeric channels could be detected on the cell surface by electrophysiological and pharmacological methods (Stewart et al., 2025). This may be due to the N458/10 epitope being inaccessible in the assembled Kv2/Kv6.4 channel in cells fixed in this manner. Indeed, we have found that antibodies to another KvS subunit (Kv5.1) have similar characteristics (Ferns et al., 2025). Consequently, we tested alternative fixation and antigen retrieval methods and identified conditions that yielded effective IF labeling of Kv6.4 in heterologous cells and tissue sections. These included fixation with glyoxal/acetic acid (Figure 1C), fixation with 4% formaldehyde followed by protease antigen retrieval (Figure 2A-C), and fixation with 2% formaldehyde followed by SDS antigen retrieval (Figure 2D).

**Figure 2.**
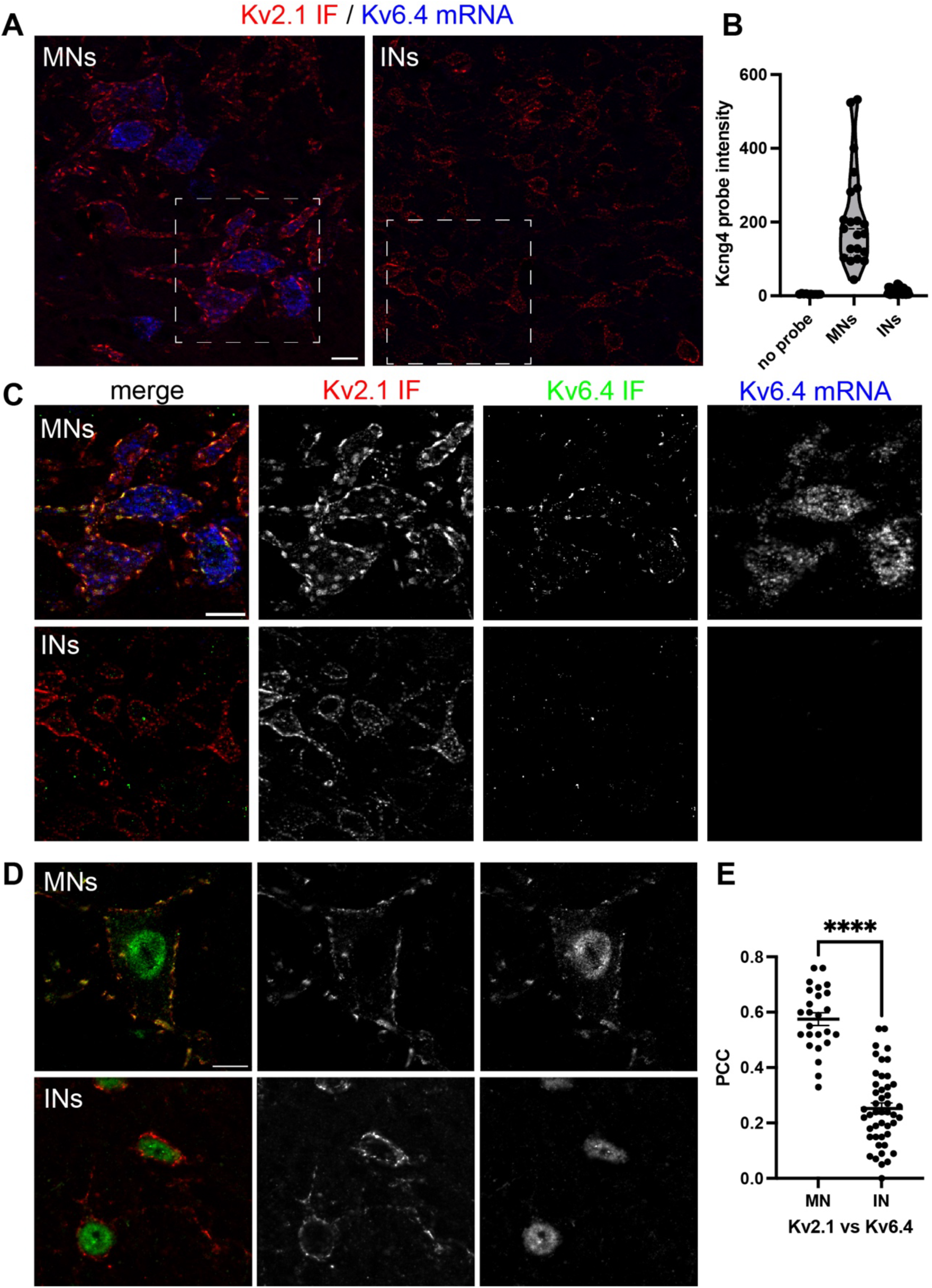
Kv6.4 transcript and protein are selectively expressed in motor neurons. (A) Cryostat sections of mouse lumbar spinal cord were hybridized using RNAscope with probes targeting Kcng4 (Kv6.4, blue) and then immunolabeled for Kv2.1 (K89/34, red). Prominent labeling for Kv6.4 transcript was evident in motor neurons in the ventral horn, but not in interneurons in other layers of spinal cord. Robust immunolabeling for Kv2.1 was detected both in motor neurons (MNs) and interneurons (INs). Images of motor neurons and interneurons were acquired with the same exposure times and were subjected to identical linear adjustments of min/max signals for display purposes. Scale bar = 20 µm. (B) Mean intensity of Kcng4 RNAscope probe labeling in Kv2.1-positive spinal cord neurons. Kcng4 probe labeling was limited to motor neurons with a broad range of intensities of probe labeling across motor neurons (n= 21 MNs and 42 INs, 2 RNAscope experiments). (C) Enlargements of boxed areas in (A) showing RNAscope using probes for Kcng4 transcript (blue) combined with immunolabeling for Kv2.1 (red) and Kv6.4 (green). Kv6.4 mRNA labeling and protein immunolabeling are detected in the same motor neurons, whereas neither are detected in interneurons. Scale bar = 20 µm. (D) Motor neuron-specific immunolabeling for Kv6.4 is also observed in samples prepared with 2% formaldehyde, pH 6 fixation and an SDS antigen-retrieval step. Representative images from n=3 spinal cords. Note that nuclear labeling for Kv6.4 (green) but not for Kv2.1 (red) occurred using this method. Scale bar = 10 µm. (E) Pearson’s correlation coefficient (PCC) values for Kv2.1 versus Kv6.4 immunolabeling are significantly higher in motor neurons compared to interneurons (unpaired t test, **** p<0.0001, n=25 MNs and 46 INs, 2 spinal cords).

To characterize Kv6.4 expression in spinal cord, we first performed RNAscope experiments on fixed, frozen sections of mouse lumbar spinal cord to localize Kv6.4 transcript at the cellular level. The sections were then immunolabeled for Kv2.1 and Kv6.4 subunits to colocalize subunit proteins. As shown in previous studies (Muennich and Fyffe, 2004; Wilson et al., 2004), we observed robust immunolabeling for Kv2.1 in spinal cord, with prominent expression in motor neurons in the ventral horn, as well as in interneurons in other layers of the cord. By contrast, labeling for Kv6.4 transcript was largely restricted to motor neurons located in both the lateral and medial motor columns (Figure 2A). Robust Kv6.4 transcript labeling was observed in both the soma and proximal dendrites of motor neurons and varied in intensity between motor neurons (Figure 2B). Much weaker Kv6.4 transcript labeling was detected in a very small number of interneurons scattered in other layers of spinal cord. Indeed, quantification of Kv6.4 mRNA labeling intensity in Kv2.1 IF-positive neurons showed that it is highly enriched in motor neurons compared to interneurons, and that labeling intensity varies between motor neurons (Figure 2A,B).

In experiments combining RNAscope with Kv2.1 and Kv6.4 immunolabeling, we found that Kv6.4 immunolabeling was prominent in motor neurons in the ventral horn, and we consistently observed co-labeling for Kv6.4 protein and transcript in individual neurons (Figure 2C). In contrast, Kv6.4 immunolabeling was largely undetectable in interneurons in other layers, even though they immunolabeled robustly for Kv2.1 (Figure 2C). A similar cellular pattern of Kv6.4 immunolabeling was observed using other fixation/antigen retrieval conditions (Figure 2D). Differential expression of Kv6.4 is also reflected in significantly higher Pearson Correlation Coefficient (PCC) values for Kv2.1 vs Kv6.4 immunolabeling in motor neurons compared to interneurons (Figure 2E). Thus, Kv6.4 transcript and protein immunolabeling are both highly enriched in motor neurons compared to interneurons. In addition, the close correspondence between Kv6.4 transcript labeling and protein immunolabeling supports the specificity of the N458/10 monoclonal antibody.

### Kv6.4 is coclustered with Kv2 subunits at C-bouton synapses

Further multiplex immunolabeling experiments revealed additional insights into Kv2/Kv6.4 channel composition and localization in motor neurons. First, we found that Kv2.1 and Kv6.4 immunolabeling colocalized extensively on the motor neuron soma and proximal dendrites, where they were detected in large micron-sized clusters (Figure 3A). Second, Kv2.1 and Kv6.4 immunolabeling intensity varied between motor neurons, as shown by the differing hues in different neurons (Figure 3A, insert). This implies that there is some heterogeneity in Kv2 and Kv6.4 subunit amounts in different motor neurons, which could impact Kv2 channel composition. Third, Kv6.4 immunolabeling also colocalized with that of Kv2.2, which is co-clustered with Kv2.1 (Figure 3B). Indeed, we found similar PCC values for Kv2.1 vs Kv6.4 and Kv2.2 vs Kv6.4 immunolabeling, which are only slightly lower than the PCC values for Kv2.1 vs Kv2.2 immunolabeling (Figure 3C). Fourth, we found that the Kv2 channel auxiliary subunit AMIGO-1 also co-clustered with Kv2 and Kv6.4 subunits in motor neurons (Figure 3D,E). This is evident in the extensive overlap of Kv2.1, Kv6.4 and AMIGO-1 labeling and in their significant PCCs (Figure 3E) and is similar to the precise colocalization of Kv2 channels and AMIGO-1 at ER-PM junctions observed in brain neurons (Bishop et al., 2018). Finally, we found that Kv2.1 and Kv6.4 co-clustered immediately beneath choline acetyltransferase (ChAT) positive synaptic terminals (Figure 3F). This is evident in intensity profile plots, where the peaks for Kv2.1 and Kv6.4 immunolabeling correspond closely and are offset from the peak for ChAT immunolabeling in the presynaptic terminal (Figure 3G). This is consistent with previous studies showing that Kv2.1 clusters in motor neurons occur at cholinergic C-bouton synapses (Muennich and Fyffe, 2004; Wilson et al., 2004). These findings identify Kv6.4 as a novel component of Kv2 channels that are clustered at C-bouton synapses in spinal motor neurons.

**Figure 3.**
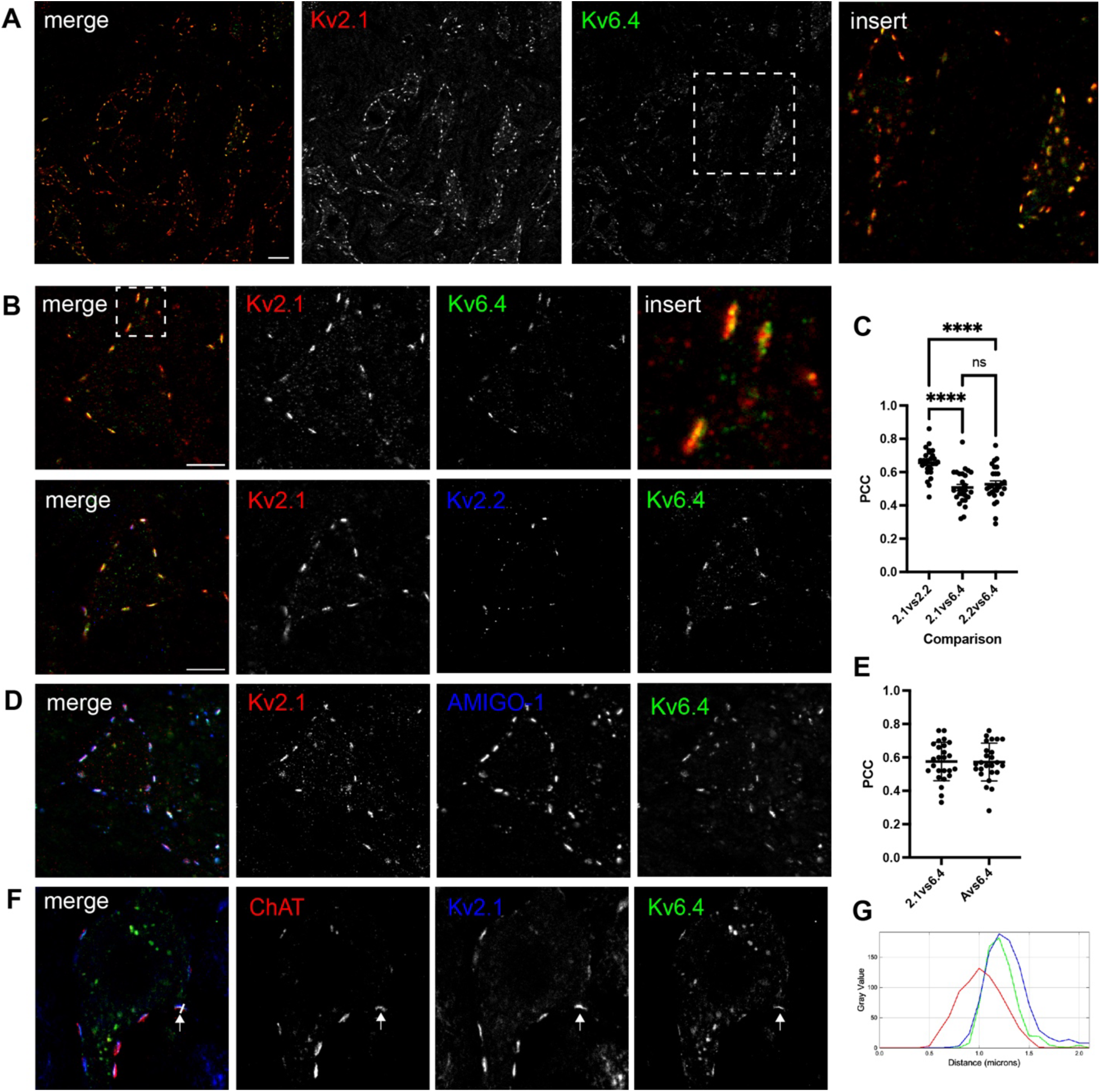
Kv6.4 is co-clustered with Kv2 channels at C-boutons in motor neurons. (A) Immunofluorescent labeling of spinal cord fixed with 9% glyoxal/8% acetic acid shows that Kv6.4 (N458/10, green) colocalizes extensively with Kv2.1 (K89/34, red) on motor neurons in the ventral horn. However, the ratio of Kv2.1 and Kv6.4 immunolabeling varies, as shown by the different hues of immunolabeling in neighboring neurons (insert). Scale bar = 20 µm. (B) Higher magnification images show that Kv6.4 (green) is co-clustered with Kv2.1 (red) and Kv2.2 (blue) on the soma and proximal dendrites of motor neurons. Scale bar = 10 µm. (C) In motor neurons, Pearson’s correlation coefficient (PCC) values are similar for Kv2.1 versus Kv6.4 and Kv2.2 versus Kv6.4 immunolabeling, indicating that Kv6.4 colocalizes with both Kv2 subunits (one way ANOVA and Tukey’s multiple comparisons test, **** p<0.0001, n=27 neurons; 2 spinal cords fixed with 9% glyoxal/8% acetic acid). (D) Immunolabeling for the AMIGO-1 auxiliary subunit (L86A/37, blue) shows that it co-localizes with Kv2.1 (K89/34, red) and Kv6.4 (N458/10, green), evident in their extensive overlap and high PCC values (panel E, n=29 neurons). (F) Immunolabeling for ChAT (pAb, red) shows that C-boutons (arrow) are juxtaposed to Kv2.1 (KC, blue) and Kv6.4 (N458/10, green) clusters on the soma of motor neurons. This is also evident in the intensity profile plot (panel G), in which the peak ChAT immunolabeling (red) is slightly offset from the peaks for Kv2.1 (blue) and Kv6.4 immunolabeling (green).

### Kv2.1 and Kv6.4 clustering at C-boutons are reduced in Kv2.1 S590A mutants

C-bouton synapses on motor neurons are also sites of close apposition between the ER and PM (*i.e.*, ER-PM junctions), and several ER and PM resident proteins are clustered at these sites. In heterologous cells and brain neurons, Kv2 channels cluster at ER-PM junctions by interacting with VAP proteins via the phospho-FFAT/PRC motif in their C-terminal domain (Johnson et al., 2018; Kirmiz et al., 2018a; Di Mattia et al., 2020). To test whether Kv2 channels localize to C-bouton associated junctions by the same mechanism, we compared Kv2 channel clustering in wild-type (WT) and Kv2.1 S590A gene-edited mice (Matsumoto et al., 2023), which lack a critical phosphorylation site in the Kv2.1 PRC domain required for binding to VAP proteins (Lim et al., 2000; Kirmiz et al., 2018a) Indeed, we observed a striking reduction in Kv2.1 clustering in motor neurons in spinal cord from Kv2.1 S590A mutant mice, with most Kv2.1 labeling being diffusely distributed on the motor neuron cell bodies and proximal dendrites (Figure 4A,B). One exception was occasional, intense Kv2.1 aggregates that localized to the proximal portion of the axon initial segment (AIS), as identified by ankyrin-G labeling (Figure 4A). This is consistent with previous work showing that Kv2.1 is localized to the AIS by a distinct molecular mechanism (Jensen et al., 2017). In contrast, Kv2.2 clusters were still observed in motor neurons from Kv2.1 S590A mutant mice (Figure 4A,B). To quantitatively compare Kv2 channel clustering, we measured coefficient of variation (CV) values for Kv2.1 and Kv2.2 immunolabeling in motor neurons. The CV value (CV: SD/mean) is used as a measure of non-uniformity of subcellular distribution, with clustered distributions having high CV values and uniform or diffuse signals having low CV values (Bishop et al., 2015; Kirmiz et al., 2018b; Kirmiz et al., 2018a). Notably, CV values for Kv2.1 were significantly decreased in motor neurons in Kv2.1 S590A mutant mice as compared to those in WT mice, whereas CV values for Kv2.2 were not significantly altered (Figure 4C). Consistent with this, the PCCs for Kv2.1 versus Kv2.2 immunolabeling in motor neurons were significantly decreased in Kv2.1 S590A mutant mice compared to WT mice (Figure 4C).

**Figure 4.**
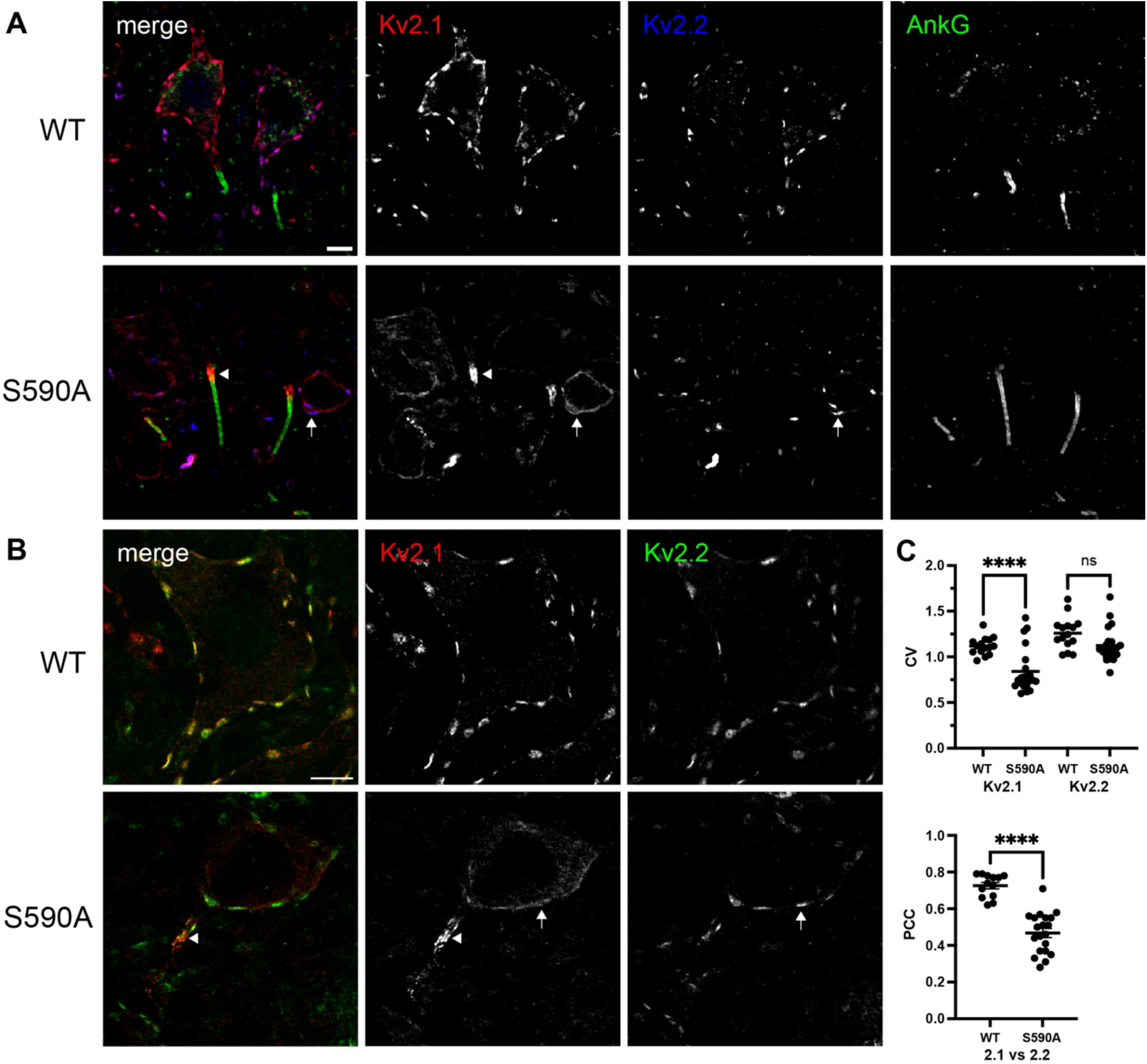
Kv2.1 channel clustering is severely reduced in Kv2.1 S590A gene-edited mice. (A) Immunolabeling for Kv2.1 (K89/34, red), Kv2.2 (2.2C, blue) and Ankyrin G (N106/36, green) in motor neurons of wild-type (WT) and Kv2.1 S590A gene-edited mice, in which Kv2.1 lacks the ability to bind VAP proteins. In Kv2.1 S590A mice, Kv2.1 immunolabeling is largely diffusely distributed on the motor neuron soma (arrow), but Kv2.2 clusters remain (arrow). In addition, prominent Kv2.1 clusters (arrowhead) are detected at the proximal portion of the axon initial segment (AIS), identified by Ankyrin G immunolabeling (green). Scale bar = 10 µm. (B) Higher magnification images showing the diffuse distribution of Kv2.1 (K89/34, red, arrow) except at the start of the AIS (arrowhead), and clustered distribution (arrow) of Kv2.2 (2.2C, green) in Kv2.1 S590A knock-in motor neurons. Scale bar = 10 µm. (C) Clustering of Kv2.1 and Kv2.2 were compared in motor neurons of WT and Kv2.1 S590A mice by measuring the coefficient of variation of labeling intensity (CV: SD/mean of pixel intensity). CV values for Kv2.1 were reduced in Kv2.1 S590A mice, whereas CV values for Kv2.2 were not significantly different (one way ANOVA and Tukey’s multiple comparisons test, **** p<0.0001, n=15 WT and 22 S590A motor neurons; 2 experiments using spinal cords fixed with 2% formaldehyde, pH 6). PCC values for Kv2.1 versus Kv2.2 immunolabeling were also significantly decreased in Kv2.1 S590A motor neurons compared to WT (one way ANOVA and Tukey’s multiple comparisons test, **** p<0.0001, n=13 WT and 20 S590A motor neurons).

Similarly, we compared Kv6.4 and AMIGO-1 clustering in motor neurons of WT and Kv2.1 S590A mutants, using 9% glyoxal/8% acetic acid-fixed tissue to optimize Kv6.4 immunolabeling. We found that Kv6.4 and AMIGO-1 clustering were reduced in Kv2.1 S590A mutants, although to a lesser extent than for Kv2.1 (Figure 5A,B). The smaller decrease in CV values for AMIGO-1 and Kv6.4 compared to Kv2.1 is likely due to the subpopulation associated with Kv2.2, which remains clustered on the soma and proximal dendrites of motor neurons (Figure 4). The PCCs for Kv2.1 versus Kv6.4, and AMIGO-1 versus Kv6.4 immunolabeling were also decreased in Kv2.1 S590A mutant motor neurons compared to those in WT mice, and the size of AMIGO-1 clusters was decreased 37% in Kv2.1 S590A mutant motor neurons (Figure 5B).

**Figure 5.**
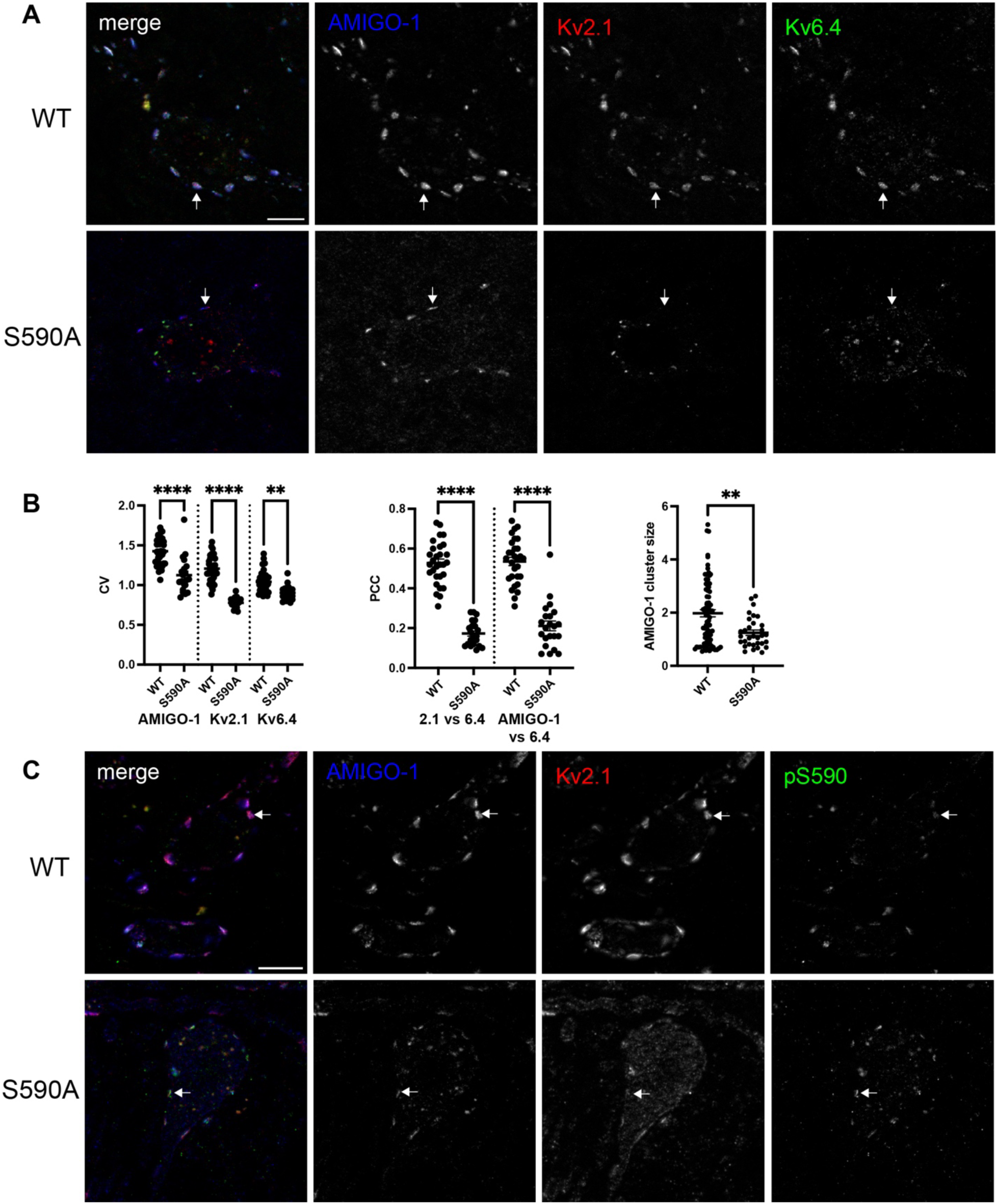
Kv2.1/Kv6.4 clustering requires S590 phosphorylation. (A) Immunolabeling for AMIGO-1 (L86A/37, blue), Kv2.1 (K89/34, red), and Kv6.4 (N458/10, green) shows that clustering of each is greatly reduced in motor neurons of Kv2.1 S590A gene-edited mice compared to WT mice (arrows). Images were acquired with the same exposure times and were subjected to identical linear adjustments of min/max signals for display purposes. Scale bar = 10 µm. (B) CV values for AMIGO-1, Kv2.1 and Kv6.4, as well as their pairwise PCC values, were all decreased significantly in Kv2.1 S590A mice compared to WT mice, reflecting their more diffuse distributions (one way ANOVA and Sidak’s multiple comparisons test, **** p<0.0001, ** p<0.005, n=39 WT and 20 S590A motor neurons; 2 experiments using spinal cords fixed with 9% glyoxal/8% acetic acid). AMIGO-1 cluster size was also decreased in the Kv2.1 S590A mutants, consistent with the de-clustering of Kv2.1 channels (n=78 WT and 32 S590A motor neurons). (C) Immunolabeling with a S590 phospho-specific antibody (L100/1, green) is detected at clusters of AMIGO-1 (L86A/37, blue) and Kv2.1 (KC, red) in WT motor neurons, but is greatly reduced at the residual AMIGO-1 clusters in Kv2.1 S590A motor neurons (arrow). Scale bar = 10 µm.

Lastly, using a phosphospecific antibody (L100/1, Table 1) specific for Kv2.1 phosphorylated at serine 590 (pS590) within the PRC domain (Cobb et al., 2015), we observed punctate labeling that colocalized with Kv2 clusters in motor neurons of WT mice, but greatly diminished labeling in those of Kv2.1 S590A mice (Figure 5C). Some residual pS590 immunolabeling that colocalized with AMIGO-1 clusters remained, possibly due to binding to the homologous phosphorylated serine (pS605) within the PRC domain of Kv2.2. This provides further evidence that the FFAT motif is phosphorylated in Kv2.1 and Kv2.2 channels clustered at C-boutons. Together, these findings indicate that Kv2.1 channel clustering at C-bouton synapses on motor neurons requires the phospho-FFAT/PRC motif and likely involves interactions with VAP proteins as in brain neurons. Moreover, as Kv6.4 subunits lack a PRC motif, Kv6.4 clustering in motor neurons is consistent with Kv6.4 co-assembly with Kv2 α subunits that contain PRC clustering domains.

### Kv6.4 and AMIGO-1 immunolabeling are reduced in Kv2 KO mice

In brain, the abundance and clustering of the AMIGO-1 auxiliary subunit at neuronal ER-PM junctions depends on its association with Kv2 channels (Bishop et al., 2018). Consistent with this, we found that AMIGO-1 immunolabeling in spinal cord was reduced in Kv2.1 and Kv2.2 KO mice. Notably, AMIGO-1 immunolabeling at motor neuron C-boutons was greatly reduced in Kv2.1 KO mice, and more moderately reduced in Kv2.2 KO mice, as compared to WT mice (Figure 6A). This distinction is also apparent in the mean intensity of AMIGO-1 labeling at somatic clusters, which is reduced by ∼60% in Kv2.1 KO, and ∼16% in Kv2.2 KO motor neurons (Figure 6B). These findings suggest that the abundance and clustering of AMIGO-1 in motor neurons depends on Kv2 channels, as it does in brain neurons. In addition, the 3.75-fold greater reduction of AMIGO-1 labeling in Kv2.1 KO compared to Kv2.2 KO mice suggests that motor neurons express significantly more Kv2.1 than Kv2.2 subunits, as shown previously for mouse brain (Bishop et al., 2018).

**Figure 6.**
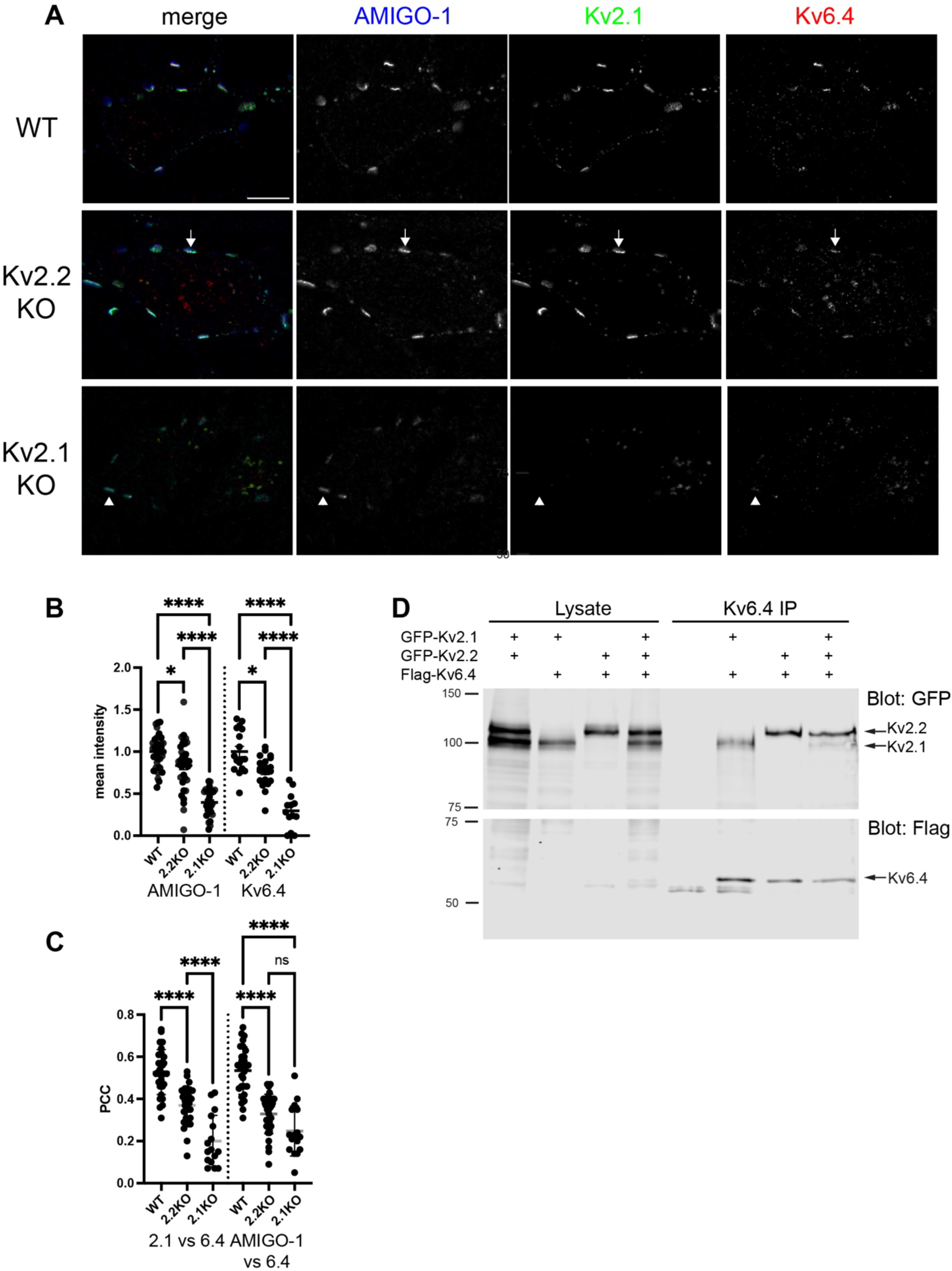
Kv6.4 immunolabeling is significantly reduced in Kv2.1 KO motor neurons. (A) Immunolabeling of motor neurons for AMIGO-1 (L86A/37, blue), Kv2.1 (K89/34, green) and Kv6.4 (N458/10, red) in WT, Kv2.2 KO, and Kv2.1 KO spinal cord. The intensity of both AMIGO-1 and Kv6.4 clusters is severely reduced in Kv2.1 KO motor neurons and modestly reduced in Kv2.2 KO motor neurons. Images were acquired with the same exposure times and were subjected to identical linear adjustments of min/max signals for display purposes. Scale bar = 10 µm. (B) Mean intensity of AMIGO-1 and Kv6.4 immunolabeling of motor neurons in WT, Kv2.2 KO, and Kv2.1 KO spinal cord. Compared to WT, AMIGO-1 immunolabeling is reduced by 16% in Kv2.2 KO and 60% in Kv2.1 KO spinal cord, and Kv6.4 immunolabeling is reduced by 23% in Kv2.2 KO and 70% in Kv2.1 KO spinal cord (one way ANOVA and Sidak’s multiple comparisons test, **** p<0.0001, * p<0.02, points represent individual neurons). (C) PCC values for Kv2.1 versus Kv6.4, and AMIGO-1 versus Kv6.4 immunolabeling are reduced more in motor neurons in Kv2.1 KO mice than in Kv2.2 KO mice (one way ANOVA and Sidak’s multiple comparisons test, **** p<0.0001, points represent individual neurons). (D) Lysates were prepared from HEK cells transfected with GFP-Kv2.1, GFP-Kv2.2, and Flag-Kv6.4 as designated in the lane labels, and IPs performed using anti-Kv6.4 mAb (N458/10). Samples of the starting lysate and IP products were size fractionated on SDS gels and immunoblotted with anti-GFP (top) and anti-Flag antibodies (bottom). GFP tagged Kv2.1 and Kv2.2 were both co-IPed with Kv6.4, consistent with their co-assembly.

In brain, the expression and clustering of another KvS subunit family member, Kv5.1, also depends on Kv2 subunits (Ferns et al., 2025). Consequently, we tested how Kv6.4 immunolabeling in motor neurons is impacted in Kv2.1 and Kv2.2 subunit KO mice. Similar to our findings for AMIGO-1, we found that Kv6.4 immunolabeling was greatly reduced in Kv2.1 KO samples, where it was only occasionally detected at Kv2.2 clusters that remained on the soma and proximal dendrites of motor neurons (Figure 6A). Kv6.4 immunolabeling was also reduced in Kv2.2 KO mice, but to a lesser extent (Figure 6A). This is supported by the mean intensity of Kv6.4 immunolabeling (Figure 6B), and PCCs for AMIGO-1 and Kv2.1 versus Kv6.4 immunolabeling (Figure 6C), which were reduced more severely in Kv2.1 KO as compared to Kv2.2 KO mice. Thus, Kv6.4 clustering is more dependent on Kv2.1 than Kv2.2 subunits, consistent with our evidence from AMIGO-1 immunolabeling that motor neurons express more Kv2.1 than Kv2.2 subunits. Kv6.4 immunolabeling was also reduced in motor neurons of Kv2.1 KO mice that were fixed with 2% formaldehyde, pH 6 (Supplemental Figure 1).

To further investigate Kv2/Kv6.4 subunit interactions, we co-expressed Kv6.4 together with Kv2.1 and/or Kv2.2 subunits in heterologous cells (Figure 6D). When expressed individually with Kv6.4, we found that Kv2.1 and Kv2.2 were each robustly co-immunoprecipitated together with Kv6.4 from cell lysates, consistent with their co-assembly into heteromeric channels. Together, our findings are most consistent with Kv6.4 co-assembly with both Kv2.1 and Kv2.2 subunits to form heteromeric Kv2/Kv6.4 channels that localize to C-boutons.

### Kv2 channels are dispensable for C-bouton and sigma-1 receptor localization

In addition to regulating motor neuron excitability and firing (Romer et al., 2019; Deardorff et al., 2021), it is speculated that Kv2 channels play a structural role at C-bouton junctions in motor neurons (Smith et al., 2024), similar to their role at ER-PM junctions in brain neurons. C-bouton ER-PM junctions in motor neurons differ from ER-PM junctions in brain neurons, however, in that they occur at synaptic sites (Deardorff et al., 2021). Consequently, we tested whether Kv2 channels are required for synaptic or ER-PM junction components at C-bouton junctions.

First, using ChAT immunolabeling to identify cholinergic nerve terminals, we found a similar pattern of C-boutons contacting motor neuron cell bodies and proximal dendrites in WT and Kv2.1/Kv2.2 DKO mice (Figure 7A). Indeed, the number of ChAT puncta per motor neuron was slightly increased and ChAT puncta size was unchanged in Kv2.1/2.2 DKO mice compared to WT (Figure 7B). Thus, C-bouton synapses on motor neurons are not dependent on Kv2 channels.

**Figure 7.**
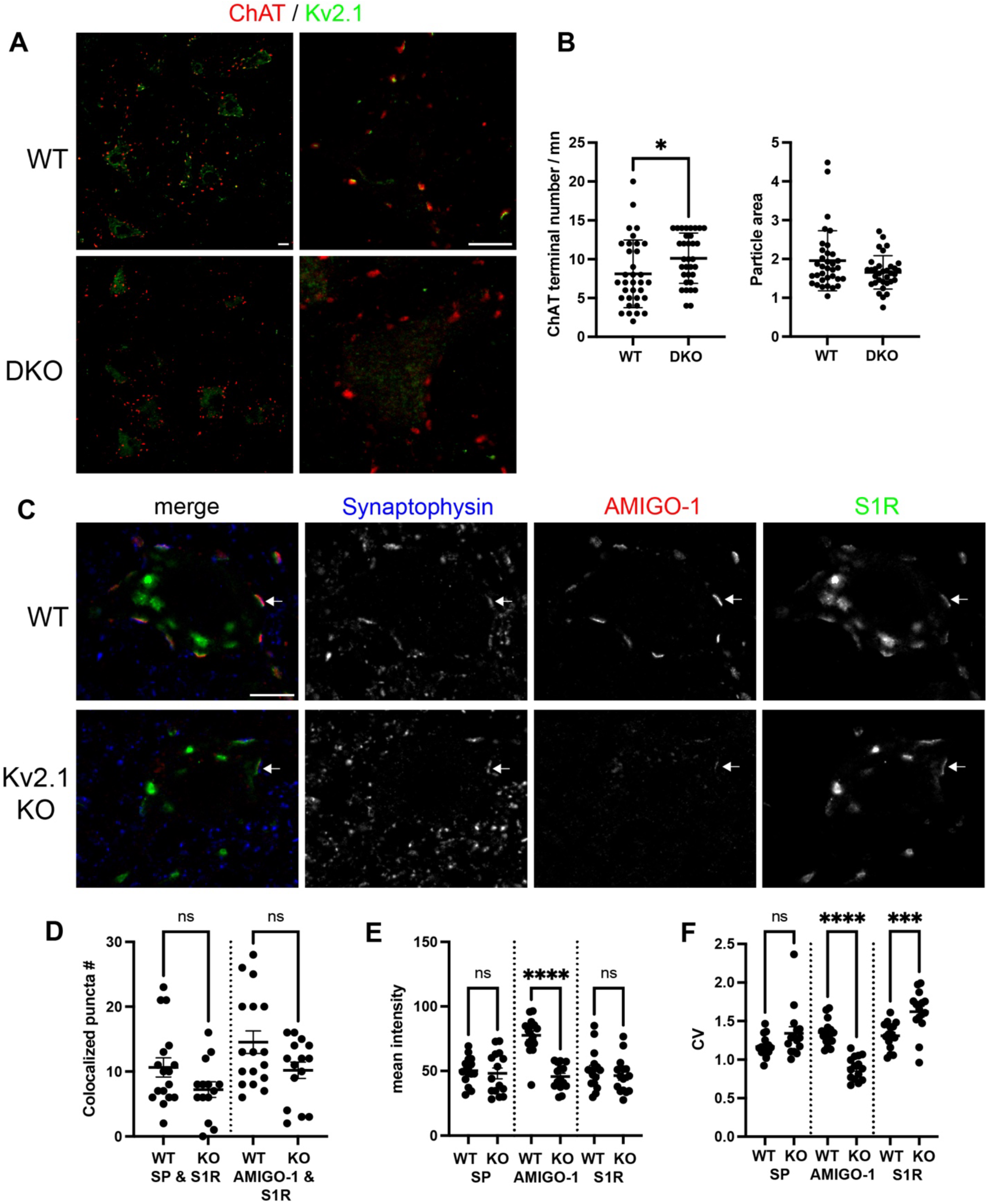
ChAT and S1R clusters are preserved in Kv2 KO motor neurons. (A) Immunolabeling for ChAT (1E6, red) and Kv2.1 (K89/34, green) reveals similar distributions of C-boutons on motor neurons from WT and Kv2.1/Kv2.2 double KO (DKO) mice. Scale bar = 20 µm. (B) ChAT puncta number is increased slightly, while ChAT puncta size is unchanged in Kv2.1/Kv2.2 DKO compared to WT motor neurons (unpaired t test, * p<0.05, n=33 WT and DKO neurons, 2 experiments using spinal cords fixed with 4% formaldehyde, pH 7.4). (C) Immunolabeling for synaptophysin (EP1098Y, blue), AMIGO-1 (L86A/37, red), and sigma-1 receptor (S1R, B-5, green) in WT and Kv2.1 KO spinal cord fixed with 9% glyoxal/8% acetic acid. S1R remains clustered near the plasma membrane in Kv2.1 KO motor neurons, where it colocalizes with weak AMIGO-1 labeling and synaptophysin-labeled terminals (arrow). Additional, cytoplasmic S1R clusters (green) are evident in both WT and Kv2.1KO neurons. Scale bar = 10 µm. (D) The number of colocalized synaptophysin (SP) and S1R puncta, and AMIGO-1 and S1R puncta are similar in WT and Kv2.1 KO motor neurons, consistent with C-bouton junctions being maintained. (E) Mean intensity of plasma membrane-associated immunolabeling for SP, AMIGO-1, and S1R in WT and Kv2.1 KO motor neurons. While the mean intensity of AMIGO-1 labeling is decreased in Kv2.1 KO compared to WT motor neurons, the intensities of SP and S1R labeling are unchanged (one way ANOVA and Sidak’s multiple comparisons test, **** p<0.0001, points represent individual neurons). (F) Similarly, CV values for AMIGO-1 labeling are decreased in Kv2.1KO compared to WT motor neurons, whereas CV values for S1R are slightly increased (one way ANOVA and Sidak’s multiple comparisons test, **** p<0.0001, *** p<0.001, points represent individual neurons).

Second, we compared the localization of the sigma-1 receptor (S1R) in motor neurons of WT and Kv2.1 KO mice. The S1R is an ER-resident protein that is clustered beneath C-boutons, where it may regulate ion channel trafficking and activity, as well as ER-mitochondrial Ca^2+^ exchange (Lievens and Maurice, 2021). Moreover, mutations in S1R cause early onset ALS (Luty et al., 2010; Al-Saif et al., 2011), whereas S1R activating ligands are protective in ALS disease models (Mancuso and Navarro, 2017; Lievens and Maurice, 2021). In WT motor neurons, we found that S1R immunolabeling colocalized extensively with Kv2.1 clusters (Figure 7C), although their immunolabeling was often slightly offset, consistent with S1R being clustered in the ER beneath plasma membrane Kv2.1 clusters. In Kv2.1 KO motor neurons, we found that S1R remained clustered beneath nerve terminals, together with remaining AMIGO-1 and Kv2.2 (Figure 7C). Moreover, the number of colocalized synaptophysin and S1R puncta (Figure 7D), mean intensity of S1R immunolabeling (Figure 7E), and S1R clustering near the plasma membrane (Figure 7F) were not reduced in motor neurons of Kv2.1 KO compared to WT mice. A normal pattern of S1R clustering in motor neurons was also observed in Kv2.2 KO mice. Thus, Kv2 channels are dispensable for S1R clustering at C-boutons.

## 4. Discussion

In general, KvS subunit family members have restricted regional and cellular mRNA expression patterns compared to the more broadly expressed Kv2.1 and Kv2.2 subunits. Consequently, the co-assembly of KvS and Kv2 subunits may create diverse, neuron-specific Kv2 channel subtypes, which are tailored to the varying functional requirements of different neuronal populations. A lack of immunological and pharmacological tools has limited the study of KvS subunits, however, and the composition and localization of native Kv2/KvS channel proteins remains largely unknown. Here, we show that KvS subunit Kv6.4 is specifically expressed in spinal motor neurons, and co-clusters with Kv2.1 and Kv2.2 subunits at C-bouton associated ER-PM junctions. Kv6.4 clustering is dependent on Kv2.1 and Kv2.2 subunits, consistent with their co-assembly into heteromeric channels. As Kv6.4 subunits confer distinct properties to Kv2 channels, inclusion of Kv6.4 may tailor Kv2 channel function to the unique functional needs of somatic motor neurons.

### Kv2/Kv6.4 channel composition and localization at C-bouton junctions

As recently shown (Smith et al., 2024; Stewart et al., 2024), we find that both Kv2.1 and Kv2.2 subunit proteins are expressed in motor neurons and clustered beneath C-boutons. As the abundance of the AMIGO-1 auxiliary subunit depends on its association with Kv2.1 and Kv2.2 subunits in brain (Bishop et al., 2018), we compared AMIGO-1 abundance at C-boutons in WT, Kv2.1 KO and Kv2.2 KO motor neurons. Notably, AMIGO-1 levels were decreased more severely in Kv2.1 KO (∼60%) compared to Kv2.2 KO motor neurons (∼16%), suggesting that Kv2.1 subunits predominate over Kv2.2 subunits by almost 4-fold. This is similar to the estimated ∼5-fold higher expression of Kv2.1 compared to Kv2.2 that was reported in mouse brain based on similar considerations (Bishop et al., 2018). Our findings that AMIGO-1 colocalizes with and tracks abundance of Kv2 channels in motor neurons also suggest that its physiological roles (Peltola et al., 2016) likely impact all Kv2 channel subtypes at C-boutons.

We find that the electrically silent Kv6.4 subunit is also expressed in motor neurons at both the transcript and protein levels. Our findings are consistent with *in situ* hybridization and transcriptomic studies available in public databases but also reveal that Kv6.4 expression is highly specific to motor neurons within spinal cord. Although we observe some heterogeneity in Kv6.4 transcript levels between motor neurons, less heterogeneity is evident in immunolabeling for Kv6.4 protein. Indeed, Kv6.4 is detectable in most if not all motor neurons in lumbar spinal cord. Elsewhere in the nervous system, Kv6.4 mRNA expression has been noted in several subclasses of DRG neurons (Zheng et al., 2019), as well as in motor nuclei in the brainstem (Allen Brain Atlas). Thus, its expression is largely in neurons involved in motor and sensory function.

Our findings also provide strong evidence that Kv6.4 co-assembles with Kv2 subunits to form heteromeric Kv2/Kv6.4 channels in motor neurons. First, we find that Kv6.4 is co-clustered with Kv2 subunits beneath C-boutons. Second, Kv6.4 clustering is severely reduced in Kv2.1 KO mice, and moderately reduced in Kv2.2 KO mice. This indicates that Kv6.4 expression and clustering is dependent on Kv2 subunits, and presumably their co-assembly into heteromeric channels. Third, the greater reduction in Kv2.1 KO compared to Kv2.2 KO mice, suggests that Kv6.4 is predominantly found co-assembled with Kv2.1 subunits. This is likely due to the higher expression of Kv2.1 compared to Kv2.2, as our expression studies in heterologous cells indicate that Kv6.4 co-assembles with both Kv2 subunits. Expression and clustering of Kv5.1 in cortical neurons is also dependent on Kv2 subunits, and unassembled Kv5.1 subunits are more prone to ubiquitination and degradation (Ferns et al., 2025). Thus, the dependency of KvS subunit expression on Kv2 subunits may extend to multiple KvS subunit family members. Together, these findings suggest that motor neurons express multiple Kv2 channel subtypes, which likely include Kv2.1 homomers, Kv2.1/Kv2.2 heteromers, and Kv2/Kv6.4 heteromers. The relative abundance of Kv2/Kv6.4 heteromeric channels compared to Kv2.1/Kv2.2 channels remains to be determined.

Another notable finding is that Kv2/Kv6.4 channels are specifically clustered at C-bouton junctions. Plasma membrane Kv2 channels localize to and organize ER-PM junctions by binding to ER-localized VAPA/B proteins, and this interaction is mediated by their FFAT/PRC domain and requires phosphorylation of S590 (Lim et al., 2000; Johnson et al., 2018; Kirmiz et al., 2018a). Consistent with this, we observed that Kv2.1 was largely diffusely distributed in motor neurons of Kv2.1 S590A mutant mice, indicating that Kv2 channels in motor neurons are localized by the same molecular mechanism as in brain neurons. Similarly, Kv6.4 was clustered at C-boutons in WT mice but largely de-clustered in Kv2.1 S590A mutant mice. As Kv6.4 lacks a FFAT/PRC domain, these findings suggest that Kv6.4 is clustered at C-boutons through its association with PRC domain-containing Kv2 subunits (i.e., in heteromeric Kv2/Kv6.4 channels). It further implies that Kv2 channel clustering does not require a PRC domain in all four subunits, as Kv2/Kv6.4 heteromers would contain three or fewer PRC domains. Recently, we found that Kv2/Kv5.1 heteromeric channels in cortical neurons are also clustered at ER-PM junctions (Ferns et al., 2025). Thus, although all KvS subunit family members lack a PRC domain, they may commonly localize to ER-PM junctions through their association with Kv2 channels.

An interesting observation in Kv2.1 S590A motor neurons was the robust clustering of Kv2.1 in the proximal portion of the AIS in motor neurons. Kv2.1 channels have previously been shown to localize to this sub-compartment in WT motor neurons (Deardorff et al., 2021). This finding supports that Kv2.1 clustering at the AIS occurs by a molecular mechanism distinct from that which determines its clustering on the soma and proximal dendrites, as suggested by a previous study based on expression of a variety of Kv2.1 deletion and mutation constructs expressed in cultured hippocampal neurons (Jensen et al., 2017). That study identified an AIS localization motif (aa 720-745) located in the C-terminal tail of Kv2.1 distal to the PRC domain and showed that phosphorylation sites in this region are critical for Kv2.1 clustering at the AIS. Our findings confirm and extend these findings by showing that Kv2.1 localizes to the AIS of spinal motor neurons in vivo independent of PRC domain phosphorylation and putative VAP interactions.

### Kv2/Kv6.4 channel function at C-bouton junctions

Previous studies have reported diverse functions for Kv6.4 subunits in neuronal and non-neuronal cell types. Targeted deletion of the Kv6.4 subunit in mice causes male sterility due to disturbed spermiogenesis (Regnier et al., 2017), and mutations in Kv6.4 in zebrafish cause developmental ear defects (Jedrychowska et al., 2024). Moreover, human Kv6.4 variants have been linked to migraine (Lafreniere and Rouleau, 2012) and to reduced labor pain during childbirth (Lee et al., 2020), likely by impacting sensory neuron excitability and firing. The mechanistic basis for the effects of Kv6.4 variants in sensory neurons remain unclear and could be due to either Kv6.4 loss-of-function or dominant-negative inhibition of Kv2.1 function (Lacroix et al., 2024; Tewari et al., 2024). However, these findings support that Kv6.4 subunits play an important physiological role in specific neurons.

In motor neurons, Kv6.4 subunits have the potential to modify both the conducting and non-conducting functions of Kv2 channels at C-bouton junctions. Cholinergic C-bouton inputs on motor neurons are activated during repetitive firing associated with locomotor activity (Miles et al., 2007), and genetic inactivation or ablation of C-bouton inputs impairs increases in motor neuron firing and muscle activation (Zagoraiou et al., 2009; Nascimento et al., 2020). This enhanced firing involves Kv2 channels (Nascimento et al., 2020), which act as high threshold channels that permit shorter interspike intervals and increased repetitive firing rates in motor neurons (Romer et al., 2019), as well as cortical and hippocampal neurons (Liu and Bean, 2014; Honigsperger et al., 2017). How Kv6.4 subunits contribute to Kv2 function in motor neurons remains to be determined, however. In heterologous cells, Kv6.4 exerts a ∼40 mV hyperpolarizing shift in the voltage-dependence of Kv2.1/Kv6.4 channel inactivation (Bocksteins et al., 2012; Bocksteins et al., 2017) and consequently, the impact of Kv6.4 subunits on Kv2 currents is expected to increase with hyperpolarization. Moreover, in cultured chick motor neurons, Kv6.4 overexpression has been reported to increase firing frequency (Muller et al., 2014). Thus, Kv2/Kv6.4 channels likely contribute to C-bouton regulation of motor neuron firing, but their impact could vary with different patterns of excitation.

Another important, non-conducting function of Kv2 channels is to organize ER-PM junctions on neuronal somata and proximal dendrites (Johnson et al., 2018; Kirmiz et al., 2018b; Johnson et al., 2019; Vierra and Trimmer, 2022). This structural role is mediated by the interaction of PM Kv2.1 channels with ER-localized VAP proteins (Johnson et al., 2018; Kirmiz et al., 2018a), and helps recruit diverse proteins involved in Ca^2+^ signaling (Vierra et al., 2019), lipid handling (Kirmiz et al., 2019; Sun et al., 2019), and PKA signaling (Vierra et al., 2023). The ultrastructure and molecular composition of ER-PM junctions is heterogenous, however, and varies between neuron subtypes and even within individual neurons (Wu et al., 2017; Deardorff et al., 2021; Vierra et al., 2023; Vullhorst et al., 2023). Indeed, C-bouton-associated junctions are unique in that they occur at sites of presynaptic cholinergic innervation, and they differ significantly in their molecular composition compared to ER-PM junctions in brain neurons. Perhaps predictably, we find that Kv2 channels are dispensable for C-bouton synaptic contacts on motor neurons. Less predictably, we find that Kv2.1 channels are also dispensable for S1R clustering at the underlying ER-PM junctions. S1R is an ER protein implicated in ion channel regulation and ER-mitochondrial Ca^2+^ signaling, and its mutation causes early-onset ALS (Mancuso and Navarro, 2017; Lievens and Maurice, 2021). Kv2 channels and S1R co-localize extensively beneath C-boutons, however we found no significant reduction in its abundance or clustering adjacent to the plasma membrane in motor neurons of Kv2.1 KO mice. Preservation of S1R clusters suggests that ER-PM junctions are maintained in Kv2.1 KO motor neurons, even though Kv2.1 is the major Kv2 channel subunit at these sites. Potentially, Kv2.2 channels or pro-neuregulin which also bind ER VAP proteins (Kirmiz et al., 2018a; Vullhorst et al., 2023) may maintain ER-PM contacts in Kv2.1 KO motor neurons. Moreover, multiple proteins can form membrane contact sites and the formation of C-bouton associated junctions may not depend on any single component.

While Kv2 channels may not be required for maintenance of C-bouton junctions per se, they may regulate the composition and function of these specialized microdomains. For example, Kv2 channels may recruit as-yet-unidentified proteins to C-bouton junctions or otherwise regulate signaling at these specialized microdomains. Similarly, Kv6.4 subunits could provide additional specificity by recruiting select proteins, which are not normally localized by Kv2 homomeric channels. These questions warrant further investigation as multiple components of C-bouton junctions have been implicated in motor neuron diseases and response to injury (*i.e.*, VAPB, S1R, neuregulin/ErbB4), highlighting the importance of these unique subcellular domains as signaling hubs for motor neuron health and survival.

## Acknowledgements

This work was supported by NIH research grant R03TR004200 to MF and JTS, and R21NS101648 and R01NS114210 to JST. We thank members of the Trimmer laboratory for their support and useful discussions of the data.

