## Supplemental Figure 1 for "Kv2/Kv6.4 heteromeric potassium channels are expressed in spinal motor neurons and localized at C-bouton synapses"

**Supplemental Figure 1. Kv6.4 immunolabeling is reduced in Kv2.1 KO motor neurons**

(A) Immunolabeling of motor neurons for Kv2.1 (K89/34, red), Kv2.2 (2.2C, blue), and Kv6.4 (N458/10, green) in WT and Kv2.1 KO spinal cord fixed with 2% formaldehyde, pH 6. The intensity of Kv6.4 clusters is reduced in Kv2.1 KO motor neurons compared to WT. Images were acquired with the same exposure times and were subjected to identical linear adjustments of min/max signals for display purposes. Scale bar = 10 μm.

(B) PCC values for Kv2.1 versus Kv6.4 immunolabeling are reduced in Kv2.1KO mice (one way ANOVA and Sidak's multiple comparisons test, **** $p<0.0001$, points represent individual neurons).

(C) Motor neuron from a Kv2.1 KO mouse immunolabeled for Kv2.2 (2.2C, blue), and Kv6.4 (N458/10, green). Profile plot across line scan shows that weak Kv6.4 immunolabeling is detectable at some Kv2.2 clusters in Kv2.1 KO motor neurons.

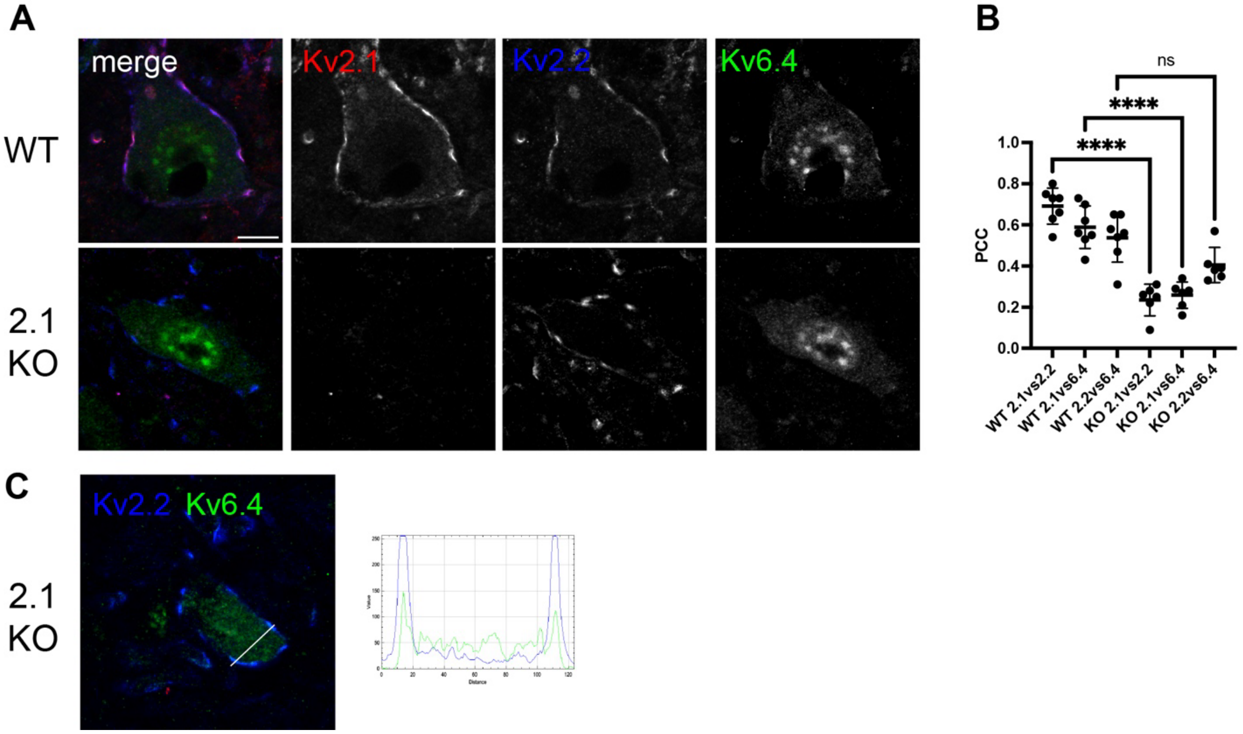
